# One-step Generation of Zebrafish Carrying a Conditional Knockout-Knockin Visible Switch via CRISPR/Cas9-Mediated Intron Targeting

**DOI:** 10.1101/827352

**Authors:** Jia Li, Hong-yu Li, Shan-ye Gu, Hua-Xing Zi, Lai Jiang, Jiu-lin Du

**Affiliations:** Institute of Neuroscience, State Key Laboratory of Neuroscience, CAS Center for Excellence in Brain Science and Intelligence Technology, Shanghai Research Center for Brain Science and Brain-Inspired Intelligence, Chinese Academy of Sciences, Shanghai 200031, China; University of Chinese Academy of Sciences, Beijing 100049, China; Department of Anesthesiology and Surgical Intensive Care Unit, Xinhua Hospital, Shanghai Jiao Tong University School of Medicine, Shanghai 200092, China; School of Life Science and Technology, ShanghaiTech University, Shanghai 200031, China

**Keywords:** NHEJ, non-HR, knockin, conditional knockout, visible switch, zCKOIS, zebrafish

## Abstract

The zebrafish has been becoming a popular vertebrate animal model in biomedical research. However, it is still challenging for making conditional gene knockout (CKO) zebrafish due to the low efficiency of homologous recombination (HR). Here we report an efficient non-HR-based method for generating zebrafish carrying a CKO and knockin (KI) switch (zCKOIS) coupled with dual-color fluorescent reporters. Using this strategy, we generated *hey2*^*zCKOIS*^ which served as a *hey2* KI reporter with EGFP expression. Upon Cre induction in targeted cells, the *hey2*^*zCKOIS*^ was switched to a non-functional CKO allele *hey2*^*zCKOIS-inv*^ associated with TagRFP expression, enabling to visualize CKO alleles. Thus, the simplification of the design, and the visibility and combination of both CKO and KI alleles’ engineering make our zCKOIS strategy an applicable CKO approach for zebrafish.

## INTRODUCTION

Knockin (KI) animals carrying exogenous sequences integrated at specific genomic loci are invaluable tools for biomedical research. To understand the role of lethal genes in post-embryonic functions, it is usual to use KI animals carrying two *loxP* insertions at interested genomic loci to generate conditional gene knockout (CKO) animals **(Yu and Bradley, 2001)**. The zebrafish (*Danio rerio*) is a vertebrate animal model excellent for in *vivo* imaging of biological events. However, to knockin two *loxP* sites in the zebrafish genome is still a big challenge due to the unavailability of gene targeting techniques for zebrafish embryonic stem cells and the low efficiency of homologous recombination (HR) in fast-developing fertilized zebrafish eggs (**Robles et al., 2011**). Although HR-mediated KI was reported to insert exogenous sequences into the genome, such as the *green fluorescent protein* (*GFP*),*loxP*, and gene-trap cassette, the HR efficiency is low **(Sugimoto, et al., 2017; Hoshijima, et al., 2016; Zu, et al., 2013)**. This low efficiency of the gene targeting via HR is a bottleneck for generating KI and CKO alleles in zebrafish. We previously developed a non-HR-mediated efficient KI strategy without destroying targeted genes, in which a Cas9 target is selected in an intron and the donor plasmid is integrated into the targeted intron **(Li, et al., 2015)**. This strategy has been used to efficiently make KI zebrafish and mouse **(Lovett-Barron, et al., 2019; Mu, et al., 2019; Suzuki, et al., 2019; Li, et al., 2015)**.

Here we develop a novel method for making <u>z</u>ebrafish <u>CKO</u> and KI <u>s</u>witch (zCKOIS) in one step based on our previously reported non-HR-mediated intron targeting, in which a floxed and invertible gene-trap cassette with an RNA slice acceptor is inserted in the intron. Although Cre-mediated switch of the gene-trap cassette was reported to make CKO zebrafish, it failed to label the wild-type cells and the knockout cells, making it impossible to distinguish which cells are the mutant **(Sugimoto, et al., 2017)**. Whereas the advantage of our design is that we can see the wild-type cells and the mutant cells directly by colors: without Cre expression, the zCKOIS cassette will not be inverted and a KI reporter with green fluorescence will be transcripted under the control of the target gene’s endogenous promoter; in the present of Cre, the cassette will be inverse, and the target gene will be destroyed, associated with the expression of a KO reporter with red fluorescence. Using our strategy, we generated a *hey2*^*zCKOIS*^ fish line and observed green fluorescence (i.e., KI reporter) in various cell types, including glial cells, endothelial cells (ECs) and haematopoietic stem cells (HSCs), similar to the expression pattern of the endogenous *hey2* in zebrafish and mice **(Rowlinson and Gering, 2010; Satow, et al., 2001; Zhong, et al., 2001)**. By injecting Cre mRNA, we then validated that the inverted *hey2*^*zCKOIS*^ (*hey2*^*zCKOIS-inv*^) coupled with red fluorescence (i.e., KO reporter) is a non-functional KO allele, because homogenous *hey2*^*zCKOIS-inv/zCKOIS-inv*^ exhibited severe defects in tail circulation of the dorsal aorta (DA), a typical phenotype of the *hey2* point mutant gridlock **(Zhong, et al., 2001; Zhong, et al., 2000)**. Finally, we achieved EC-specific KO of *hey2* by crossing the *hey2*^*zCKOIS*^ with the KI line *Ki(flk1-P2A-Cre)*, in which Cre expression is driven by the EC-specific promoter *flk1*. We found that CKO of *hey2* in ECs did not cause defects in tail circulation, suggesting that the Hey2 plays a non-EC-autonomous role in regulating circulation development.

## RESULTS

### Generation of a *hey2*^*zCKOIS*^ Zebrafish Line

We selected *hey2* to test our strategy. The Hey2, a transcriptional repressor known as an effector of Notch signaling, is broadly expressed in various cell types, including ECs, HSCs, and glial cells **(Rowlinson and Gering, 2010; Satow, et al., 2001; Zhong, et al., 2001)**. We first made a *hey2-GSG-P2A-EGFP* KI donor based on our previous study **(Li, et al., 2015)**. The glycine-serine-glycine (GSG) sequence enables efficient cleavage of the P2A **(Wang, et al., 2015)**. Then we modified the EGFP KI donor by inserting in the left arm with a gene trap cassette, which includes an invertible splice acceptor site followed by a red fluorescent protein (TagRFP) via a P2A sequence for multicistronic expression (Sugimoto, et al., 2017) (**Figure 1A**). We name it <u>z</u>ebrafish <u>C</u>ondition <u>K</u>nock<u>O</u>ut and knock<u>I</u>n <u>S</u>witch (zCKOIS), because this design is supposed to realize one-step generation of an allele with both EGFP-labeled KI and TagRFP-labeled KO. We co-injected the zCKOIS donor plasmid, a short guide RNA (sgRNA) and the mRNA of zebrafish codon-optimized Cas9 (zCas9) into one-cell-stage fertilized eggs and raised these embryos to adulthood (**Figure 1A**). Those F0 were then outcrossed to wild-type (WT) zebrafish, and their F1 progenies were screened for EGFP signal. Two F0 founders with germline transmission were identified in a total of 21 F0 fish. We then used F1 embryos from one of the founders for the following experiments. PCR of genomic DNA and sequencing analysis of the F1 progenies confirmed the inheritance of the genomic integration of the F0 founder (**Figure 1B** and **Figure S1A**). Reverse transcription (RT)-PCR and sequencing also confirmed that EGFP was expressed with the endogenous *hey2* transcripts (**Figure 1C** and **Figure S1B**). Hereafter, we refer this line as *hey2*^*zCKOIS*^.

**Figure 1.**
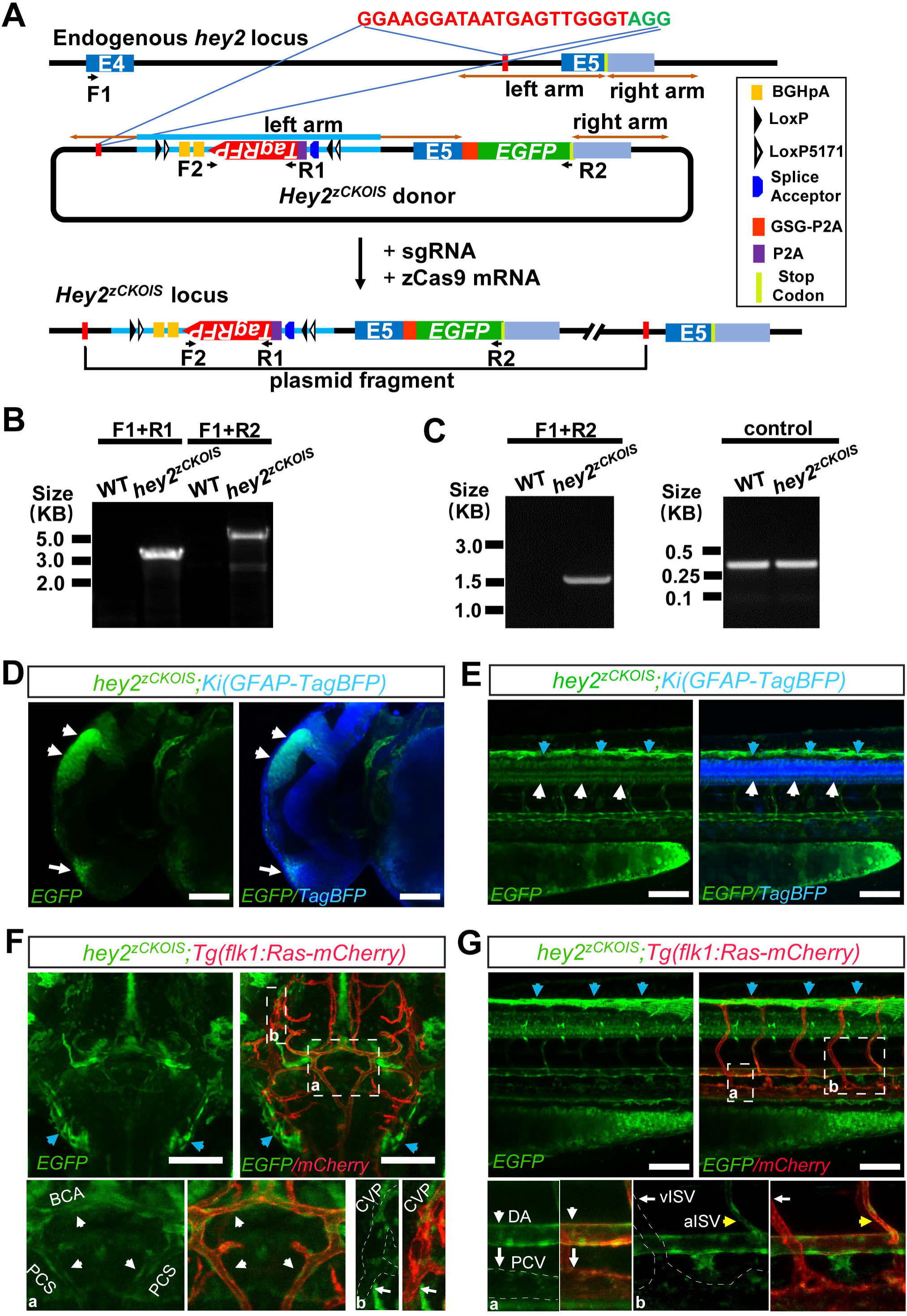
Generation of a *hey2*^*zCKOIS*^ Zebrafish Line. **(A)** Schematic of the intron targeting-mediated strategy for generating *hey2*^*zCKOIS*^ zebrafish by using the CRISPR/Cas9 system. The zebrafish *hey2* has 5 exons, and E4 and E5 represent the 4^th^ and 5^th^ exons, respectively. The *hey2*^*zCKOIS*^ donor was integrated into the *hey2* locus after co-injection of the donor with the sgRNA and zCas9 mRNA. The sgRNA target sequence is showed in red and the protospacer adjacent motif (PAM) sequence in green. The left and right arm sequences of the donor plasmids are indicated by the brown lines with double arrows. The left arm in the donor is 3300 bp in length, including the original left arm in the genome and an inverted TagRFP cassette sequence. The right arm is 1107 bp in length. GSG-P2A is glycine-serine-glycine-P2A sequence. **(B)** PCR analysis of the genomic DNA obtained from the F1 progenies of the *hey2*^*zCKOIS*^ founder. A 3.2-kb band was amplified by using the F1 and R1 primers, and a 5.3-kb band was amplified by using the F1 and R2 primers. The two bands are only present in *hey2*^*zCKOIS*^ embryos, but not in the wild-type (WT) group. The F1, R1, F2, and R2 primers are indicated in **(A). (C)** Left, RT-PCR analysis using the cDNA of *hey2*^*zCKOIS*^ F1 embryos. A 1.5-kb band amplified by the F1 and R1 primers only appears in *hey2*^*zCKOIS*^ embryos. Right, control RT-PCR using the primers binding *hey2* coding sequence showed a 0.3-kb band. **(D)** Projected confocal images (lateral view) of a *hey2*^*zCKOIS*^;*Ki(GFAP-TagBFP)* embryos at 1.5 dpf, showing EGFP expression in glial cells in the brain labeled by TagBFP. *Ki(GFAP-TagBFP)* is a KI line with TagBFP expression specific in glial cells. White arrowheads, the midbrain; Arrow, the forebrain. Scale bars: 100 μm. **(E)** Projected confocal images (lateral view) of a *Ki(GFAP-TagBFP);hey2*^*zCKOIS*^ embryo at 3.5 dpf, showing EGFP expression in the glia cells at the spinal cord (white arrow heads). Cyan arrowheads, non-specific signals on the skin. Scale bars, 100 μm. **(F and G)** Projected confocal imaging of the *hey2*^*zCKOIS*^;*Tg(flk1:Ras-mCherry)* embryos at 3.5 dpf. **(F)** EGFP expression in the basal communicating artery (BCA) and postcardinal communicating segment (PCS) in the brain (dorsal view). Top left, EGFP signal; top right, merged signals (EGFP/mCherry). The outlined areas labeled a and b in the top right panel are enlarged below. White arrowheads, BCA and PCS; White arrows, choroidal vascular plexus (CVP). **(G)** EGFP expression in the dorsal aorta (DA) but not in the postcardinal vein (PCV) in the trunk (lateral view). Arterial intersegmental vessels (aISVs) extending from the DA exhibited more EGFP signal than venous ISVs (vISVs) extending from the PCV. Top left, EGFP signal; top right, merged signals (EGFP/mCherry). The outlined areas labeled a and b in the top right panel are enlarged below. White arrowheads, DA and aISVs; White arrows, PCV and vISVs. Cyan arrowheads, non-specific signals on the skin. Scale bars: 100 μm.

Consistent with the reported expression patterns of *hey2* in zebrafish and mice **(Rowlinson and Gering, 2010; Satow, et al., 2001; Zhong, et al., 2001)**, *hey2*^*zCKOIS*^ showed intensive expression of EGFP in glial cells and vascular ECs (**Figures 1D-1G**). Interestingly, EGFP was found to express in arteries, such as the basal communicating artery (BCA) and postcardinal communicating segment (PCS) in the brain, and the dorsal aorta (DA) and arterial intersegmental vessels (aISVs) in the trunk. However, in veins, such as the choroidal vascular plexus (CVP) in the brain, and the postcardinal vein (PCV) and venous ISVs (vISVs) in the trunk, EGFP expression was hardly detected (**Figures 1F** and **1G**). This is consistent with the role of *hey2* in the fate decision of arterial ECs **(Korten, et al., 2013; Zhong, et al., 2001)**. Taking together, it indicates that the EGFP-labeled KI of *hey2*^*zCKOIS*^ recaptures the endogenous expression pattern of *hey2* in zebrafish.

### Characterization of the *hey2*^*zCKOIS*^ Allele

To characterize the *hey2*^*zCKOIS*^ allele, we injected Cre recombinase mRNA into the one-cell-stage fertilized eggs of *hey2*^*zCKOIS*^ fish and raised the injected embryos to adulthood. By outcrossing to WT fish, we identified the F1 progenies with TagRFP expression, which indicates the inversion of the *hey2*^*zCKOIS*^ allele (*hey2*^*zCKOIS-inv*^) (**Figures 2A** and **2B**). PCR and DNA sequencing of the F1 genomic DNA showed the inversion of the *hey2*^*zCKOIS*^ cassette as expected (**Figure 2C** and **Figure S2A**). RT-PCR and sequencing data also confirmed the in-frame ligation of the mRNA of the *hey2* exon4 to the P2A-TagRFP cassette (**Figure 2D** and **Figure S2B**).

**Figure 2.**
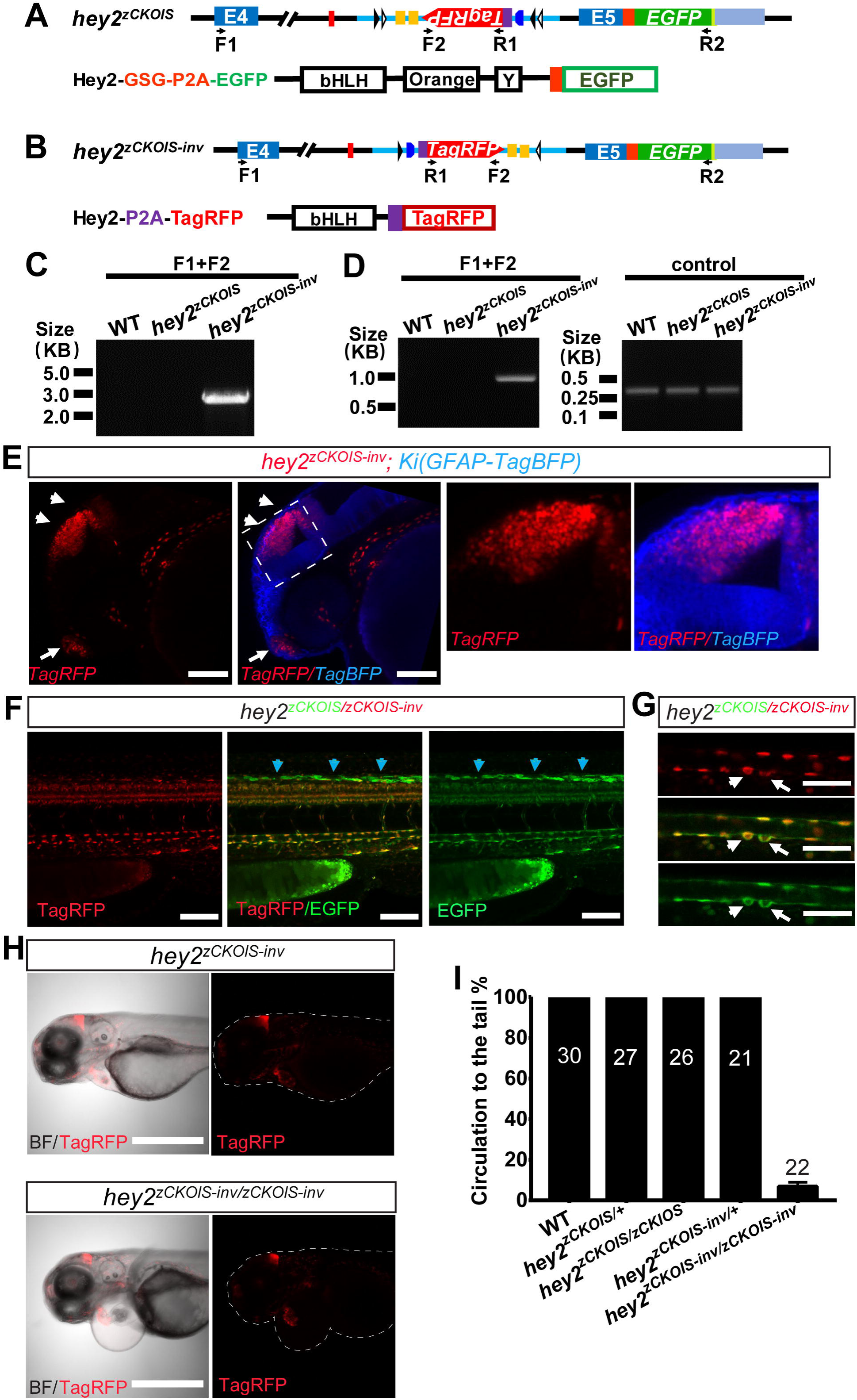
Characterization of the *hey2*^*zCKOIS-inv*^ Allele. **(A)** Schematic of the *hey2*^*zCKOIS*^ allele and the translated Hey2^zCKOIS^ protein. The endogenous Hey2 protein includes an N-terminal bHLH domain, an Orange domain, and a protein-protein interaction YRPW (“Y”) motif near the C-terminus. The Hey2^zCKOIS^ protein is a fusion of the wild-type (WT) Hey2 protein and a GSG-P2A-EGFP. **(B)** Schematic of the *hey2*^*zCKOIS-inv*^ allele produced via Cre-induced inversion of the *hey2*^*zCKOIS*^ allele. The translated Hey2^zCKOIS-inv^ protein is a truncated Hey2 with only the bHLH domain fused to a P2A-TagRFP. **(C)** PCR detection of the inversion in *hey2*^*zCKOIS-inv*^ genome. A 2.8-kb band was only present in the *hey2*^*zCKOIS-inv*^ but not in the WT or *hey2*^*zCKOIS*^ embryos. **(D)** Left, RT-PCR analysis for detection of the transcription of *hey2*^*zCKOIS-inv*^. A 0.9-kb band was amplified only in the *hey2*^*zCKOIS-inv*^, not in the WT or *hey2*^*zCKOIS*^ embryos. Right, control RT-PCR using primers binding *hey2* coding sequence showed a 0.3-kb band in all groups. **(E)** Projected confocal images (lateral view) of a *hey2*^*zCKOIS-inv*^;*Ki(GFAP-TagBFP)* embryo at 1.5 dpf, showing TagRFP expression in glia cells. The boxed area is enlarged on the right. White arrow heads, the midbrain; Arrow, the forebrain. Scale bars: 100 μm. **(F)** Projected confocal images (lateral view) of the trunk of a *hey2*^*zCKOIS/zCKOIS-inv*^ embryo at 2.5 dpf. The red fluorescence is encoded by the truncated Hey2-P2A-TagRFP in the *hey2*^*zCKOIS-inv*^ allele and the green fluorescence is encoded by the WT Hey2-GSG-P2A-EGFP in the *hey2*^*zCKOIS*^ allele. Cyan arrowheads, non-specific signals on the skin. Scale bars: 100 μm. **(G)** Confocal images of the DA in a *hey2*^*zCKOIS/zCKOIS-inv*^ embryo at 2.5 dpf showing two HSCs budding from the DA. Scale bars: 50 μm. **(H)** Bright-field images showing severe pericardial edema in the homozygous *hey2*^*zCKOIS-inv/zCKOIS-in*^ (top) but not in the heterozygous *hey2*^*zCKOIS-inv*^ (bottom) at 3.5 dpf. Left, merged images of bright filed (BF) and TagRFP. Right, TagRFP. Scale bars: 500 μm. **(I)** Percentage of embryos with different genetic backgrounds showing normal circulation to the tail. The number on the bar is the total number of 3.5-dpf embryos examined.

We then imaged *hey2*^*zCKOIS-inv*^ F1 embryos and found that, similar to the EGFP expression pattern in *hey2*^*zCKOIS*^ embryos, red fluorescence in *hey2*^*zCKOIS-inv*^ embryos was also expressed in glial cells and vascular ECs (**Figures 2E** and **2F**, and **Figures S2C** and **S2D**). Interestingly, we also observed that HSCs derived from ventral DA exhibited strong Hey2 expression during budding process (**Figure 2G**). In particular, we crossed *hey2*^*zCKOIS-inv*^ with *hey2*^*zCKOIS*^ and imaged the embryo of *hey2*^*zCKOIS/zCKOIS-inv*^. The complete overlapping of green and red signals in the embryo indicates the *hey2*^*zCKOIS-inv*^ faithfully recaptures the endogenous *hey2* expression as the *hey2*^*zCKOIS*^ allele, implying that Cre-mediated inversion does not affect gene expression (**Figures 2F** and **2G**, and **Figures S2C** and **S2D**). We noticed that in comparison with EGFP signal, TagRFP preferentially localized in the nuclear (**Figures 2E-2G**). The reason is probably that, the GSG spacer, which is crucial for the efficient cleavage of P2A, was added in the P2A-EGFP cassette but not the P2A-TagRFP cassette. This might cause the complete release of EGFP from Hey2, whereas the TagRFP might be partially fused with Hey2 and targeted to the nucleus where Hey2 is localized **(Jia, et al., 2007)**.

The exon5 of *hey2* contains important functional domains of the Hey2 protein **(Jia, et al., 2007)**. Cre-mediated inversion of TagRFP resulted in the skipping of the entire exon5 during mRNA processing (**Figure 2B**). Therefore, the *hey2*^*zCKOIS-inv*^ allele is supposed to be a loss-of-function of *hey2*. To validate it, we incrossed the *hey2*^*zCKOIS-inv/+*^ fish and obtained homozygous *hey2*^*zCKOIS-inv/zCKOIS-inv*^ embryos. Similar to the reported *hey2* point mutant (i.e., gridlock) and KO zebrafish **(Gibb, et al., 2018; Zhong, et al., 2000)**, these embryos exhibited pericardial edemas and loss of circulation to the tail (**Figures 2H** and **2I**). These detects were not observed in the embryo of heterozygous *hey2*^*zCKOIS-inv/+*^, heterozygous *hey2*^*zCKOISv/+*^ or homozygous *hey2*^*zCKOIS/zCKOIS*^ (**Figure 2I**), indicating that the *hey2*^*zCKOIS*^ is a functional KI allele and the *hey2*^*zCKOIS-inv*^ is a non-functional KO allele.

### EC-Specific KO of *hey2*

Vascular ECs are important for supporting the normal function of the circulation system. Our data above showed that *hey2* was strongly expressed in the DA and the *hey2*^*zCKOIS-inv/zCKOIS-inv*^ exhibited gridlock phenotypes in the DA characterized by the loss of circulation to the tail (see **Figures 1G** and **2I)**. To test whether the loss of *hey2* in ECs is sufficient to cause the gridlock phenotype, we crossed the *hey2*^*zCKOIS*^ fish to the KI line *Ki(flk1-P2A-Cre)*, in which the Cre expression is under control of the EC-specific *flk1* promoter **(Li, et al., 2015)**. By incrossing the fish of *hey2*^*zCKOIS*^;*Ki(flk1-P2A-Cre)*, we obtained the progenies with the background of *hey2*^*zCKOIS*^;*Ki(flk1-P2A-Cre)* that both copies of the allele was supposed to be inverted and TagRFP should be specifically expressed in ECs (**Figure 3A**). Genotyping of the genomic DNA from these progenies confirmed the inversion of the *hey2*^*zCKOIS*^ cassette (**Figure S3A**). RT-PCR data also confirmed the in-frame ligation of the mRNA of the *hey2* exon4 to the P2A-TagRFP cassette (**Figure 3B**). As expected, TagRFP was observed in ECs of arteries, including the DA and aISVs, but not in ECs of the siblings without Cre induction (**Figure 3C** and **Figure S3B**), indicating the specific loss of the *hey2* allele in ECs. As mentioned above, the *hey2* KO displayed gridlock phenotype (see **Figure 2I**). Therefore, we examined whether EC-specific KO of *hey2* could recapture the circulation defect and pericardial edemas. However, these phenotypes were not observed in the embryos with the genetic background of *hey2*^*zCKOIS/zCKOIS*^;*Ki(flk1-P2A-Cre)* (**Figure 3D**), suggesting that *hey2* may play a EC-independent role in regulating the circulation to the tail in zebrafish.

**Figure 3.**
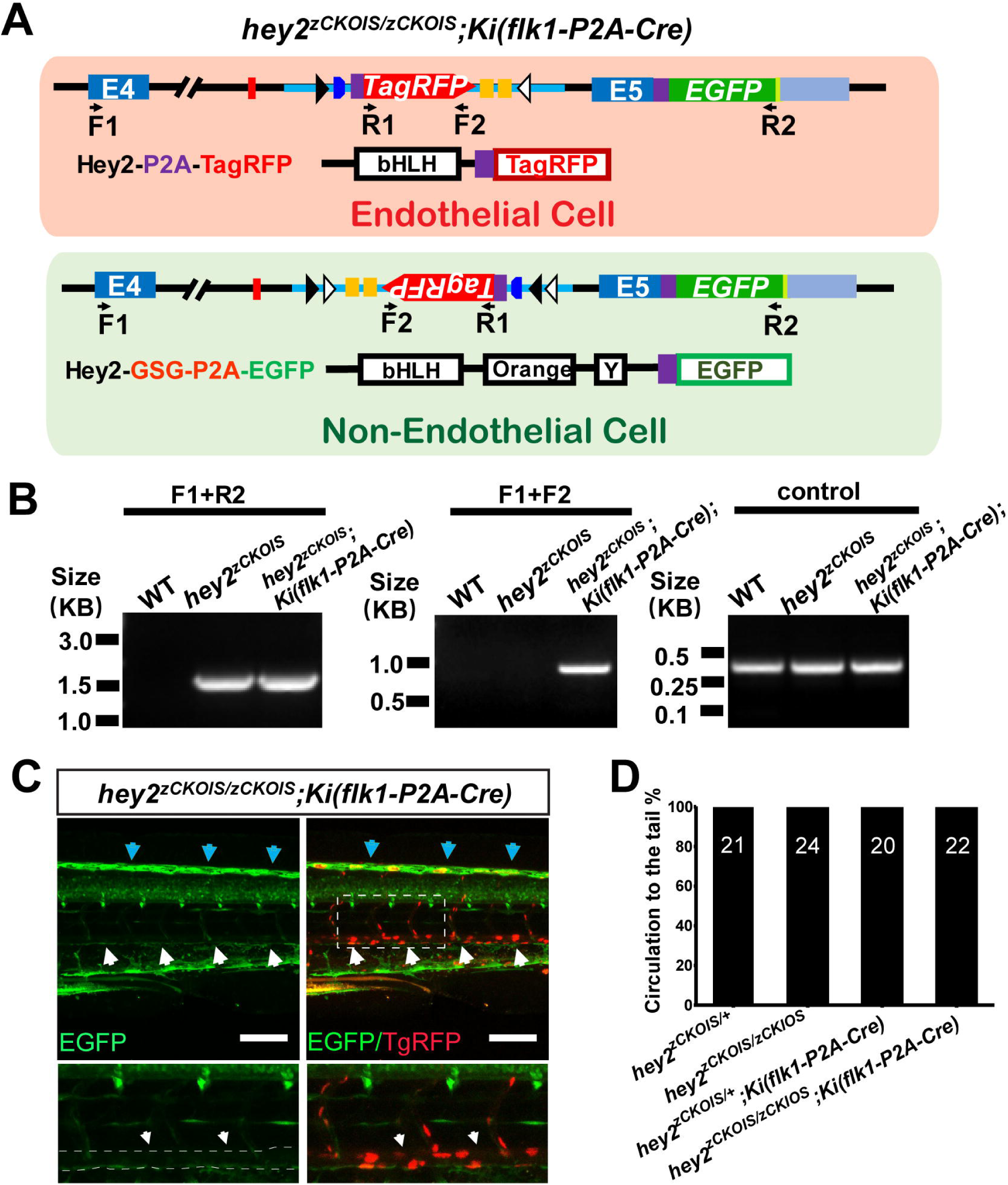
EC-Specific KO of *hey2*. **(A**) Schematic of the strategy for endothelial cell (EC)-specific KO of *hey2*. The Hey2 protein in ECs will be destroyed in the embryo carrying homozygous *hey2*^*zCKOIS*^ alleles and *Ki(flk1-P2A-Cre)* background. In non-ECs where Cre is not expressed, the *hey2*^*zCKOIS*^ will not be inverted and thus the WT Hey2 protein is expressed. **(B)** RT-PCR analysis using the cDNA for detection of the transcripts of *hey2*^*zCKOIS*^ and *hey2*^*zCKOIS-inv*^. Left, a 1.5-kb band was amplified by the F1 and R2 primers in both *hey2*^*zCKOIS*^;*Ki(flk1-P2A-Cre)* and *hey2*^*zCKOIS*^ but not in the WT embryos. Middle, a 0.9-kb band amplified by the F1 and F2 primers was only present in the *hey2*^*zCKOIS*^;*Ki(flk1-P2A-Cre)* embryos. Right, control RT-PCR using primers binding *hey2* coding sequence showed a 0.3-kb band in each group. **(C)** Projected confocal images (lateral view) of trunk vessels in a *hey2*^*zCKOIS/zCKOIS*^;*Ki(flk1-P2A-Cre)* embryos at 3.5 dpf. Cre-induced inversion of the *hey2*^*zCKOIS*^ enables the TagRFP expression in the DA and aISVs, where EGFP expression is barely deleted (arrowheads and dashed lines). Top left, EGFP signal; Top right, merged signals (EGFP/TagRFP). The outlined area is enlarged below. Cyan arrowheads, non-specific signals on the skin. Scale bars: 100 μm. **(D)** Percentage of the embryos with different genetic backgrounds showing normal circulation to the tail at 3.5dpf. The number on the bar is the total number of fish examined.

## DISCUSSION

Here we provide a simple efficient method named zCKOIS for making zebrafish lines with both visible CKO and KI alleles in one step. We showed that the zCKOIS cassette can be targeted into the last intron of *hey2* with a high efficiency via CRIPSR/Cas9 mediated non-HR insertion. As the EGFP expression in *hey2*^*zCKOIS*^ embryos is driven by the endogenous *hey2* promoter, EGFP was found to express in various cell types including glial cells and ECs. The homozygous *hey2*^*zCKOIS/zCKOIS*^ did not display any gridlock phenotype, indicating that the *hey2*^*zCKOIS*^ is a functional allele. Thus, the *hey2*^*zCKOIS*^ can be used as a *hey2* KI reporter with EGFP expression. On the other hand, by using Cre-induced inversion, we further validated that the TagRFP-tagged *hey2*^*zCKOIS-inv*^ is a non-functional allele as the homozygous *hey2*^*zCKOIS-inv/ zCKOIS-inv*^ larvae showed the obvious gridlock phenotype. Therefore, the *hey2*^*zCKOIS/zCKOIS*^ fish with a tissue specific Cre expression can realize the CKO of the *hey2* gene. Importantly, as the green color and red color represent the functional KI and non-functional KO alleles, respectively, we can easily distinguish KO cells and WT cells under microscopy just by eyes without sacrificing the fish for PCR and sequencing. This will greatly increase the screening efficiency and expand the way we detect KO events *in vivo*. For examples, we can make the inversion of the zCKOIS allele in some of the cells by using a line with weak Cre expression. Subsequently, cells with green (i.e., KI) or red color (i.e., KO) can be traced individually for exploring the distinct behaviors or the cell fate in a real-time manner *in vivo*.

For the successful one-step generation of zCKOIS of an interested gene, there are several key steps needed to be considered. The first one is how to choose a proper intron for the zCKOIS donor plasmid insertion. The ideal site for targeting is the intron just before the exon with important functional domains. The insertion of the *hey2*^*zCKOIS*^ donor into the last intron of *hey2* is because the last exon encodes important domains of the Hey2 protein, and thus the inversion of the DNA cassette in the *hey2*^*zCKOIS*^ allele will result in the loss of the last exon during *hey2* mRNA processing, guaranteeing the *hey2*^*zCKOIS-inv*^ is a non-functional KO allele. If the functional domain of an interested gene is present in the X^th^ exon, the (X-1)^th^ intron should be the suitable site for targeting. Second, it is important to select a sgRNA with a high cleavage efficiency in the intron. This is the crucial step for the successful insertion of the zCKOIS donor plasmid, because the highly efficient concurrent cleavage of the donor and target genomic sites will increase the germline transmission rate. For the case of the *hey2*^*zCKOIS*^, the cleavage efficiency of the sgRNA used was about 50%, resulting in successful identification of two founders among the total of 21 F0 fish. The third important step is the design of a zCKOIS donor. It is of importance to label the functional KI cassette and the KO cassette with different fluorescent colors, which will make it convenient to distinguish between the two genetic backgrounds by eyes. Last but not least, to achieve complete CKO in a specific cell type or tissue, the Cre expression needs to be highly specific. Leaky expression of Cre will potentially cause false-positive results. KI lines harboring the endogenous *cis*-regulatory elements of promoter/enhancer outperforms conventional transgenic lines in guaranteeing the specific expression **(Li, et al., 2015)**. Therefore, we used the Ki(*flk1-P2A-Cre*) line rather than the promoter-driven *Tg*(*flk1:Cre)* transgenic line for the inversion of the zCKOIS allele in ECs. Unexpectedly, in the *hey2*^*zCKOIS/zCKOIS*^;*Ki(flk1-P2A-Cre)*, we did not observe the gridlock phenotype found in *hey* KO and gridlock mutants, suggesting that the *hey2* of ECs is not important for regulation of the circulation in the tail of zebrafish. A previous study showed that the gridlock phenotype can be suppressed by up-regulating vascular endothelial growth factor (VEGF) or activating the VEGF pathway (Peterson, et al., 2004). Therefore, it is possible that Hey2 promotes VEGF production in the surrounding cells. Then VEGF can be secreted into extracellular space and distributes around the DA to activate VEGF downstream signals in ECs for regulating the circulation to tail. Cell type-specific Cre mediated CKO with use of the *hey2*^*zCKOIS*^ line may help to find the detailed mechanisms in the future.

In summary, the method we introduced here realizes the generation of CKO and KI reporter line in one step via efficient non-HR-mediated insertion. The simplification and combination of CKO and KI make the zCKOIS strategy an applicable approach for zebrafish and even other organisms.

## EXPERMENTIAL PROCEDURES

### Zebrafish Husbandry

Adult zebrafish were maintained in the National Zebrafish Resources of China (Shanghai, China) with an automatic fish housing system at 28°C. Embryos were raised under a 14h-10h light-dark cycle in 10% Hank’s solution that consisted of (in mM): 140 NaCl, 5.4 KCl, 0.25 Na_2_HPO_4_, 0.44 KH_2_PO_4_, 1.3 CaCl_2_, 1.0 MgSO_4_ and 4.2 NaHCO_3_ (pH 7.2). Zebrafish handling procedures were approved by the Institute of Neuroscience, Chinese Academy of Sciences.

### Production of zCas9 mRNA, sgRNAs, and Cre mRNA

The zCas9 expression plasmid pGH-T7-zCas9 (Liu, et al., 2014) was linearized by *Xba I* and used as a template for Cas9 mRNA *in vitro* synthesis and purification with the mMACHINE T7 Ultra kit (Ambion). The coding sequence of Cre was amplified from Cre plasmid by using a T7-promoter included primers. The Cre mRNA was synthesized by the same T7 Ultra kit. The sequence of sgRNAs was designed according to previously reported criteria (Chang, et al., 2013). We used the CRISPR/Cas9 design tool (*http://crispor.tefor.net/crispor.py*) to select specific targets to minimize off-target effects. The sequences of designed sgRNAs are as follows:

*hey2*: GGAAGGATAATGGTTGGGT (forward strand);

*flk1*: TCTGGTTTGGAAGGACACAG (forward strand);

*gfap*: GTGCGCAACACATAGCACCA (reverse strand);

A pair of oligonucleotides containing the sgRNA targeting sequence were annealed and cloned downstream of the T7 promoter in the PT7-sgRNA vector. The sgRNA was synthesized by the MAXIscript T7 Kit (Ambion) and purified by using the mirVana™ miRNA Isolation Kit (Ambion).

### Generation of the *hey2*^*CKOIS*^ Zebrafish Line

The *GSG-P2A-EGFP* fragment was ligated into the PMD-19-T by T-A cloning to form the *T-GSG-P2A-EGFP* vector. The left and right arms for *hey2* were amplified by the KOD-PLUS Neo DNA polymerase from WT zebrafish genomic DNA. Then, the arms were ligated to the 5’ and 3’ regions of the *GSG-P2A-EGFP* fragment in the *T-GSG-P2A-EGFP* vector, respectively. The inverted cassette with *loxP* sites, splice acceptor, TagRFP and BGHpA was amplified from the *pZwitch+1* plasmid. The resulting 1.8-kb DNA fragment was ligated to the left arm sequence via the ClaI restriction enzyme site to get the *hey2*^*CKOIS*^ donor plasmid. Finally, the zCas9 mRNA, *hey2* sgRNA, and *hey2*^*CKOIS*^ donor plasmid were co-injected into one-cell-stage fertilized zebrafish eggs. Each embryo was injected with 1 nl of solution containing 800 ng/μl zCas9 mRNA, 80 ng/μl sgRNA, and 15 ng/μl donor plasmid. To screen *hey2*^*CKOIS*^ founders, adult fish were crossed with AB wild-type (WT) zebrafish, the genomic DNA was extracted at 3 days post-fertilization (dpf), and the germline transmission was detected by PCR and imaging.

The left arm and right arm sequences of the donor plasmid were amplified from genomic DNA isolated from adult WT AB zebrafish using the following primers:

1. Left arm amplification primers: Forward: 5′- CGAGGTACCCACTCGTCGACAAAACTAGGG -3′ and Reverse: 5′- CGAGGATCCAAACGCTCCCACTTCAGTTC -3′.
2. Right arm amplification primers: Forward: 5′- CGAACCGGTTAAATGTTGGATTTAAATGT -3′ and Reverse: 5′- CGACTGCAGTAGGGTTTTAGCAGGCACCG -3′.

### RNA Preparation and First-Strand cDNA Synthesis

Total RNAs of zebrafish embryos of 1.5 dpf and 3.5dpf were extracted by using TRIzol reagent according to the manufacturer’s instructions (Invitrogen, 15596018). The extracted total RNA was used to generate the first-strand cDNA by using PrimeScript™ RT Master Mix (Takara, RR036A).

### Genotyping of *hey2*^*zCKOIS*^ and *hey2*^*zCKOIS-inv*^ Genomic Loci and RT-PCR Analysis of the Transcripts

The following primers were used for PCR of genome DNA and RT-PCR of cDNA in

**Figures 1B** and **1C, Figures 2C** and **2D, Figure 3B**, and **Figure S3A**.

F1: 5′- GATCTGCCAAGTTGGAGAAAGC -3′

F2: 5′-TCAATTAAGTTTGTGCCCCAGT -3′

R1: 5′-CACCGTGAACAACCACCACT -3′

R2: 5′-CTTGTACAGCTCGTCCATGCC -3′

*hey2* cDNA control RT-PCR primers:

F: 5′-ATGAAGCGGCCCTGTGAGGA -3′ and

R: 5′-CTTTTCCTCCTGTGGCCTGAA -3′

### Generation of the KI Line *Ki(flk1-P2A-Cre)* with Cre Expression Specific in Vascular Endothelial Cells

The *Ki(flk1-P2A-Cre)* line was generated with CRISPR/Cas9-mediated KI method as previously reported **(Li, et al., 2015)**. Briefly, the *flk1-P2A-Cre* plasmid donor was made by replacing the EGFP sequence in the flk1-P2A-EGFP donor with Cre. 1 nl of solution containing 800 pg zCas9 mRNA, 80 pg *flk1* gRNA and 15 pg *flk1* donor plasmid, was injected into zebrafish embryos at one-cell stage. The embryos were raised to adulthood for founder screening. To screen *Ki(flk1-P2A-Cre)* founders, adult fish were crossed with AB WT zebrafish, the genomic DNA was extracted at 1 dpf, and the germline transmission was detected by PCR. The founder was then crossed to the lineage tracing line *Tg(bactin2:loxP-STOP-loxP-DsRed-express)*^*sd5*^ to confirm the Cre-mediated *loxP* site deletion in vessels (**Figure S3C**) **(Bertrand, et al., 2010)**.

### Generation of the Glial Reporter KI Line *Ki(GFAP-TagBFP)*

The *Ki(GFAP-TagBFP)* line was generated as previously reported **(Li, et al., 2015)**. Briefly, The *GFAP-TagBFP* donor was made by replacing the EGFP sequence in the GFAP-EGFP donor with TagBFP. 1 nl of solution containing 800 pg zCas9 mRNA, 80 pg *gfap* gRNA and 15 pg donor plasmid, was injected into zebrafish embryos at one-cell stage. The embryos were raised to adulthood for founder screening. To screen *Ki(GFAP-TagBFP)* founders, adult fish were crossed with WT zebrafish, the genomic DNA was extracted at 1 dpf, and the germline transmission was detected by PCR and imaging.

### Confocal Imaging

Z-stack images were taken at room temperature, under an Apo LWD 25X water-immersion objective (N.A., 1.1) by using a Nikon FN1 confocal microscope (Nikon, Japan). The z-step of images ranged from 3 μm - 5 μm. To detect pericardial edemas, a Plan Fluor 10x Water objective lens (N.A.0.3) was taken with a z-step of 5 μm. The resolution of all the images was either 1024 × 1024 pixels or 512 × 512 pixels. Raw images were processed with Triangle method (ImageJ) for adjusting threshold.

## Supporting information

Figure S1. Sequencing Analysis of the Genome and Transcript of hey2zCKOIS. (Related to Figure 1)

Figure S2. Genotyping and Expression of hey2zCKOIS. (Related to Figure 2)

Figure S3. Genotyping of hey2zCKOIS;Ki(flk1-P2A-Cre) and Projected Confocal Images of hey2zCKOIS/zCKOIS and Ki(flk1-P2A-Cre). (Related to Figure 3)

## SUPPLEMENTAL INFORMATION

Supplemental Information includes three figures.

## AUTHOR CONTRIBUTIONS

J.L. and J.L.D. conceived the project and designed the experiments. J.L., H.Y.L. and S.Y.G. performed research and analyzed data. H.X.Z. made the *Ki(flk1-P2A-Cre)* zebrafish line. J.L., H.Y.L. and J.L.D. wrote the paper.

## ACKNOWLEDGEMENTS

We are grateful to Drs. N Lawson for providing the *Tg(flk1:EGFP)* line, D Traver for providing the *Tg(bactin2:loxP-STOP-loxP-DsRedEx)* line and K Kikuchi for providing the pZwitch+1 plasmid. This work was supported by the Young Scientists Fund of the National Natural Science Foundation of China (Grant No.31500849), Shanghai Municipal Science and Technology Major Project (18JC1410100, 2018SHZDZX05), the Key Research Program of Frontier Sciences (QYZDY-SSW-SMC028), the Strategic Priority Research Program (XDB32010200) of Chinese Academy of Science, the International Partnership Program, Bureau of International Co-operation of Chinese Academy of Science(153D31KYSB20170059), China Wan-Ren Program, and Shanghai Leading Scientist Program.

## REFERENCES

Bertrand, J.Y., Chi, N.C., Santoso, B., Teng, S., Stainier, D.Y., and Traver, D. (2010). Haematopoietic stem cells derive directly from aortic endothelium during development. Nature 464, 108–11.

Chang, N., Sun, C., Gao, L., Zhu, D., Xu, X., Zhu, X., Xiong, J.W., and Xi, J.J. (2013). Genome editing with RNA-guided Cas9 nuclease in zebrafish embryos. Cell research 23, 465–72.

Gibb, N., Lazic, S., Yuan, X., Deshwar, A.R., Leslie, M., Wilson, M.D., and Scott, I.C. (2018). Hey2 regulates the size of the cardiac progenitor pool during vertebrate heart development. Development 145.

Hoshijima, K., Jurynec, M.J., and Grunwald, D.J. (2016). Precise Editing of the Zebrafish Genome Made Simple and Efficient. Dev Cell 36, 654–67.

Jia, H., King, I.N., Chopra, S.S., Wan, H., Ni, T.T., Jiang, C., Guan, X., Wells, S., Srivastava, D., and Zhong, T.P. (2007). Vertebrate heart growth is regulated by functional antagonism between Gridlock and Gata5. Proc Natl Acad Sci U S A 104, 14008–13.

Korten, S., Brunssen, C., Poitz, D.M., Grossklaus, S., Brux, M., Schnittler, H.J., Strasser, R.H., Bornstein, S.R., Morawietz, H., and Goettsch, W. (2013). Impact of Hey2 and COUP-TFII on genes involved in arteriovenous differentiation in primary human arterial and venous endothelial cells. Basic Res Cardiol 108, 362.

Li, J., Zhang, B.B., Ren, Y.G., Gu, S.Y., Xiang, Y.H., and Du, J.L. (2015). Intron targeting-mediated and endogenous gene integrity-maintaining knockin in zebrafish using the CRISPR/Cas9 system. Cell Res 25, 634–7.

Liu, D., Wang, Z., Xiao, A., Zhang, Y., Li, W., Zu, Y., Yao, S., Lin, S., and Zhang, B. (2014). Efficient gene targeting in zebrafish mediated by a zebrafish-codon-optimized cas9 and evaluation of off-targeting effect. Journal of genetics and genomics = Yi chuan xue bao 41, 43–6.

Lovett-Barron, M., Chen, R., Bradbury, S., Andalman, A.S., Wagle, M., Guo, S., and Deisseroth, K. (2019). Multiple overlapping hypothalamus-brainstem circuits drive rapid threat avoidance. bioRxiv.

Mu, Y., Bennett, D.V., Rubinov, M., Narayan, S., Yang, C.T., Tanimoto, M., Mensh, B.D., Looger, L.L., and Ahrens, M.B. (2019). Glia Accumulate Evidence that Actions Are Futile and Suppress Unsuccessful Behavior. Cell 178, 27–43 e19.

Peterson, R.T., Shaw, S.Y., Peterson, T.A., Milan, D.J., Zhong, T.P., Schreiber, S.L., MacRae, C.A., and Fishman, M.C. (2004). Chemical suppression of a genetic mutation in a zebrafish model of aortic coarctation. Nat Biotechnol 22, 595–9.

Rowlinson, J.M., and Gering, M. (2010). Hey2 acts upstream of Notch in hematopoietic stem cell specification in zebrafish embryos. Blood 116, 2046–56.

Satow, T., Bae, S.K., Inoue, T., Inoue, C., Miyoshi, G., Tomita, K., Bessho, Y., Hashimoto, N., and Kageyama, R. (2001). The basic helix-loop-helix gene hesr2 promotes gliogenesis in mouse retina. J Neurosci 21, 1265–73.

Sugimoto, K., Hui, S.P., Sheng, D.Z., and Kikuchi, K. (2017). Dissection of zebrafish shha function using site-specific targeting with a Cre-dependent genetic switch. Elife 6.

Suzuki, K., Yamamoto, M., Hernandez-Benitez, R., Li, Z., Wei, C., Soligalla, R.D., Aizawa, E., Hatanaka, F., Kurita, M., Reddy, P., et al. (2019). Precise in vivo genome editing via single homology arm donor mediated intron-targeting gene integration for genetic disease correction. Cell Res 29, 804–819.

Wang, Y., Wang, F., Wang, R., Zhao, P., and Xia, Q. (2015). 2A self-cleaving peptide-based multi-gene expression system in the silkworm Bombyx mori. Sci Rep 5, 16273.

Yu, Y., and Bradley, A. (2001). Engineering chromosomal rearrangements in mice. Nat Rev Genet 2, 780–90.

Zhong, T.P., Childs, S., Leu, J.P., and Fishman, M.C. (2001). Gridlock signalling pathway fashions the first embryonic artery. Nature 414, 216–20.

Zhong, T.P., Rosenberg, M., Mohideen, M.A., Weinstein, B., and Fishman, M.C. (2000). gridlock, an HLH gene required for assembly of the aorta in zebrafish. Science 287, 1820–4.

Zu, Y., Tong, X., Wang, Z., Liu, D., Pan, R., Li, Z., Hu, Y., Luo, Z., Huang, P., Wu, Q., et al. (2013). TALEN-mediated precise genome modification by homologous recombination in zebrafish. Nat Methods 10, 329–31.

