## SUPPLEMENTAL FIGURE LEGENDS for "One-step Generation of Zebrafish Carrying a Conditional Knockout-Knockin Visible Switch via CRISPR/Cas9-Mediated Intron Targeting"

**Figure S1. Sequencing Analysis of the Genome and Transcript of *hey2^zCKOIS^*. (Related to Figure 1)**

**(A)** 5’ junction sequence of The F1 progeny of a *hey2^zCKOIS^* founder at the donor integration site. The PAM and sgRNA target sequence are shown in green and red, respectively. There is an 894 bp deletion in the intron near the sgRNA target site where the *hey2^zCKOIS^* donor plasmid was integrated. **(B)** cDNA sequence of the *hey2^zCKOIS^* transcript showing that EGFP is in-frame ligated to the exon5 of *hey2.*

**Figure S2. Genotyping and Expression of *hey2^zCKOIS^*. (Related to Figure 2)**

**(A**) 5’ junction sequences of the F1 progeny of a *hey2^zCKOIS-inv^* founder. The PAM and sgRNA target sequences are shown in green and red, respectively. There is the same 894 bp deletion in the intron near the sgRNA target site as in the *hey2^zCKOIS^*. **(B)** cDNA sequence of the *hey2^zCKOIS-inv^* transcript showing that TagRFP is in-frame ligated to the exon4 of *hey2*. **(C)** Co-localization. Scale bars, 100 μm. **(D)** Co-localization of EGFP (encoded by *hey2^zCKOIS^*) and TagRFP (encoded by *hey2^zCKOIS-inv^*) in the opercular artery (ORA) of a *hey2^zCKOIS^/^zCKOIS-inv^* embryo at 2.5 dpf. Scale bars, 50 μm.

**Figure S3. Genotyping of *hey2^zCKOIS^;Ki(flk1-P2A-Cre)* and Projected Confocal Images of *hey2^zCKOIS/zCKOIS^* and *Ki(flk1-P2A-Cre)*. (Related to Figure 3)**

**(A)** Left, PCR analysis using the genomic DNA, showing a 3.2-kb band in the *hey2^zCKOIS^* and *hey2^zCKOIS^;Ki(flk1-P2A-Cre)* groups but not in the WT group. Right, a 2.8-kb band was only present in the *hey2^zCKOIS^;Ki(flk1-P2A-Cre)* and the positive control *hey2^zCKOIS-inv^* groups, but not in the WT or *hey2^zCKOIS^* groups. **(B)** In the absent of Cre, EGFP was expressed in the DA (arrowheads), and no red fluorescent signal was not detected in the TagRFP channel. Asterisks, non-specific signals on the yolk sac. Cyan arrowheads, non-specific signals on the skin. Scale bars, 100 μm. **(C)** Projected confocal images of trunk vessels in a 3.5-dpf embryo of *Ki(flk1-P2A-Cre);Tg(bactin2:loxP-STOP-loxP-DsRedEx);Tg(flk1:EGFP).* Red, *bactin2:DsRedEx*. EGFP, *flk1:EGFP*. Scale bars, 100 μm.
