## Supplementary figures and images for "One-step Generation of Zebrafish Carrying a Conditional Knockout-Knockin Visible Switch via CRISPR/Cas9-Mediated Intron Targeting"

### Figure S1. Sequencing Analysis of the Genome and Transcript of hey2zCKOIS. (Related to Figure 1)

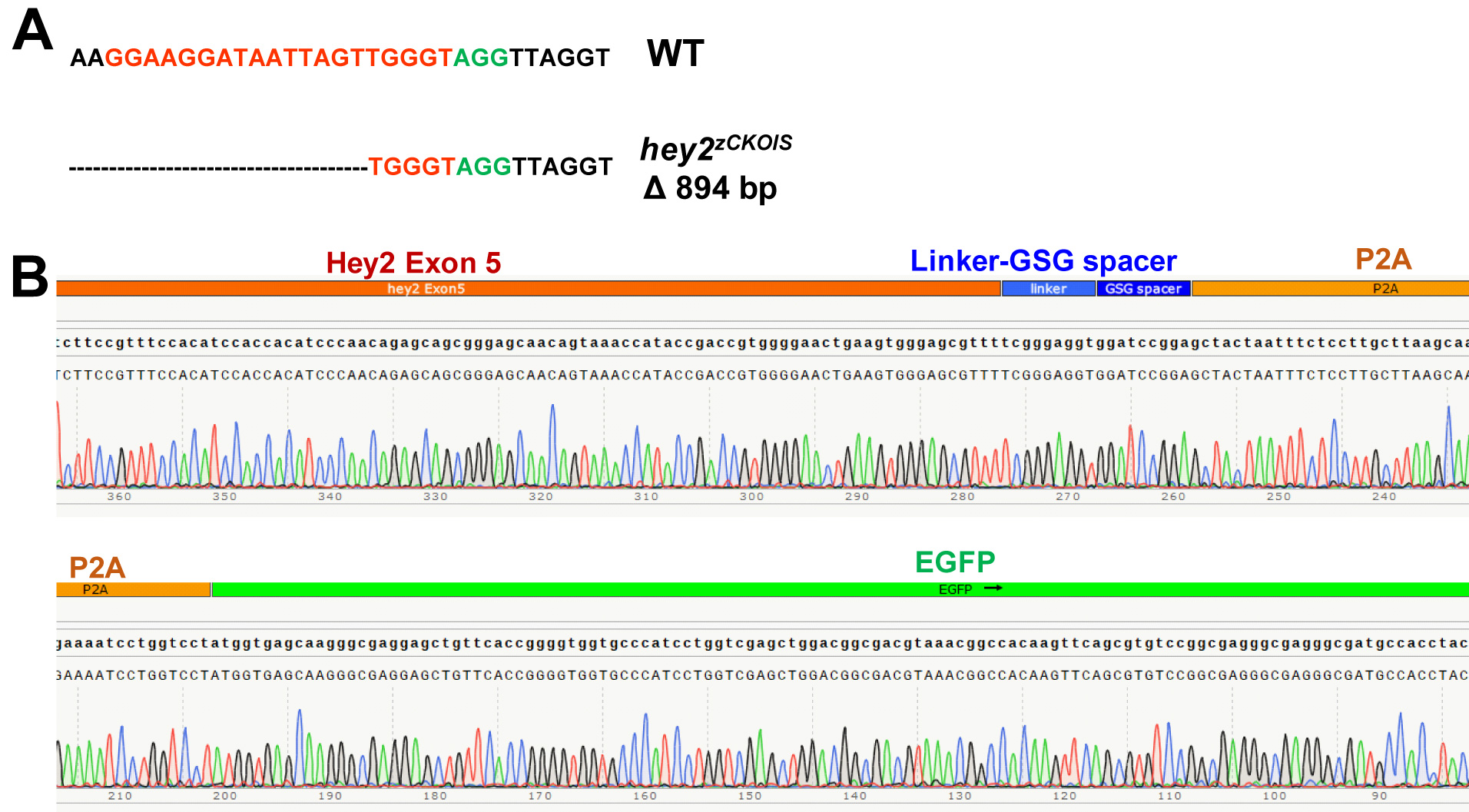

### Figure S2. Genotyping and Expression of hey2zCKOIS. (Related to Figure 2)

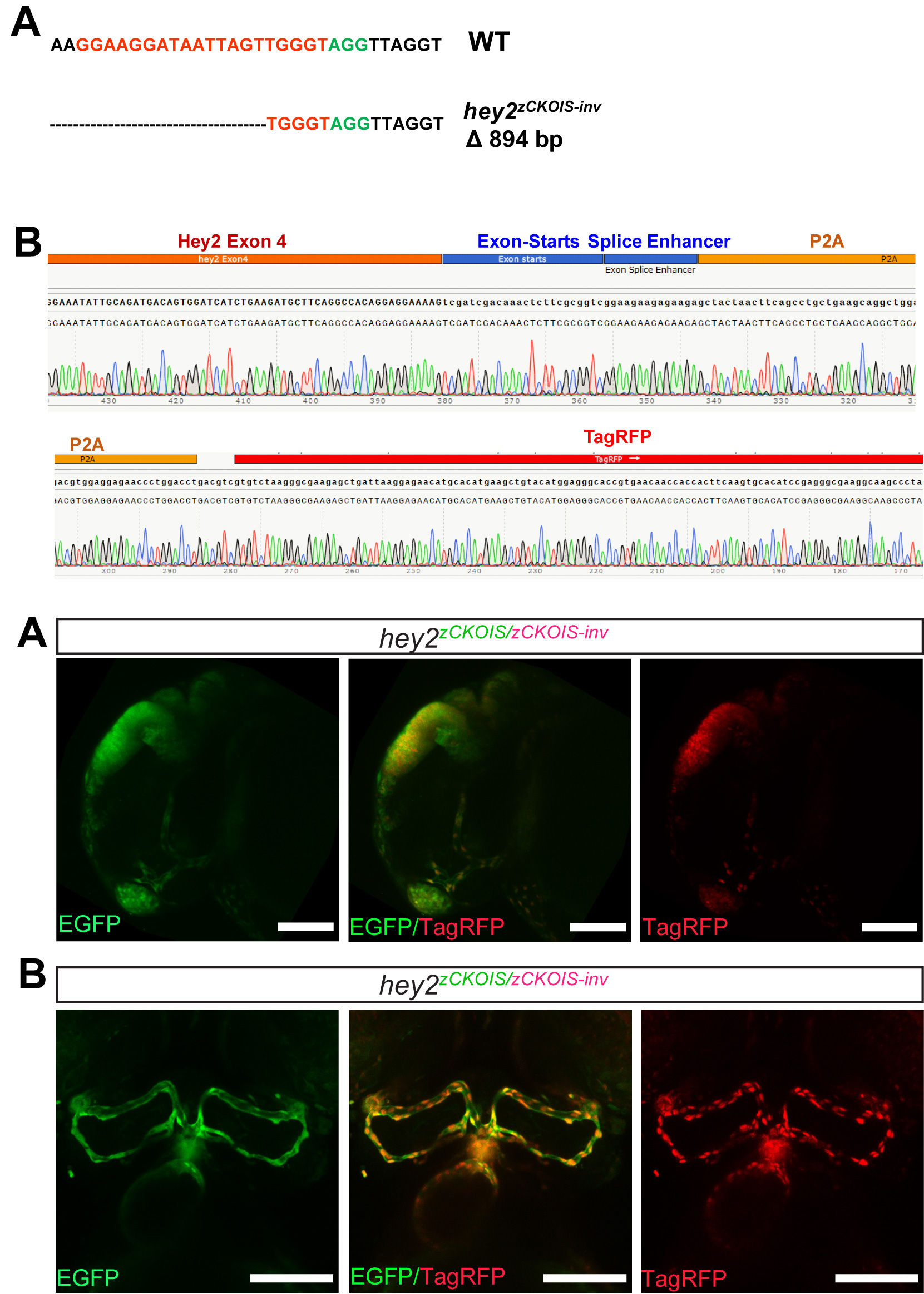

### Figure S3. Genotyping of hey2zCKOIS;Ki(flk1-P2A-Cre) and Projected Confocal Images of hey2zCKOIS/zCKOIS and Ki(flk1-P2A-Cre). (Related to Figure 3)

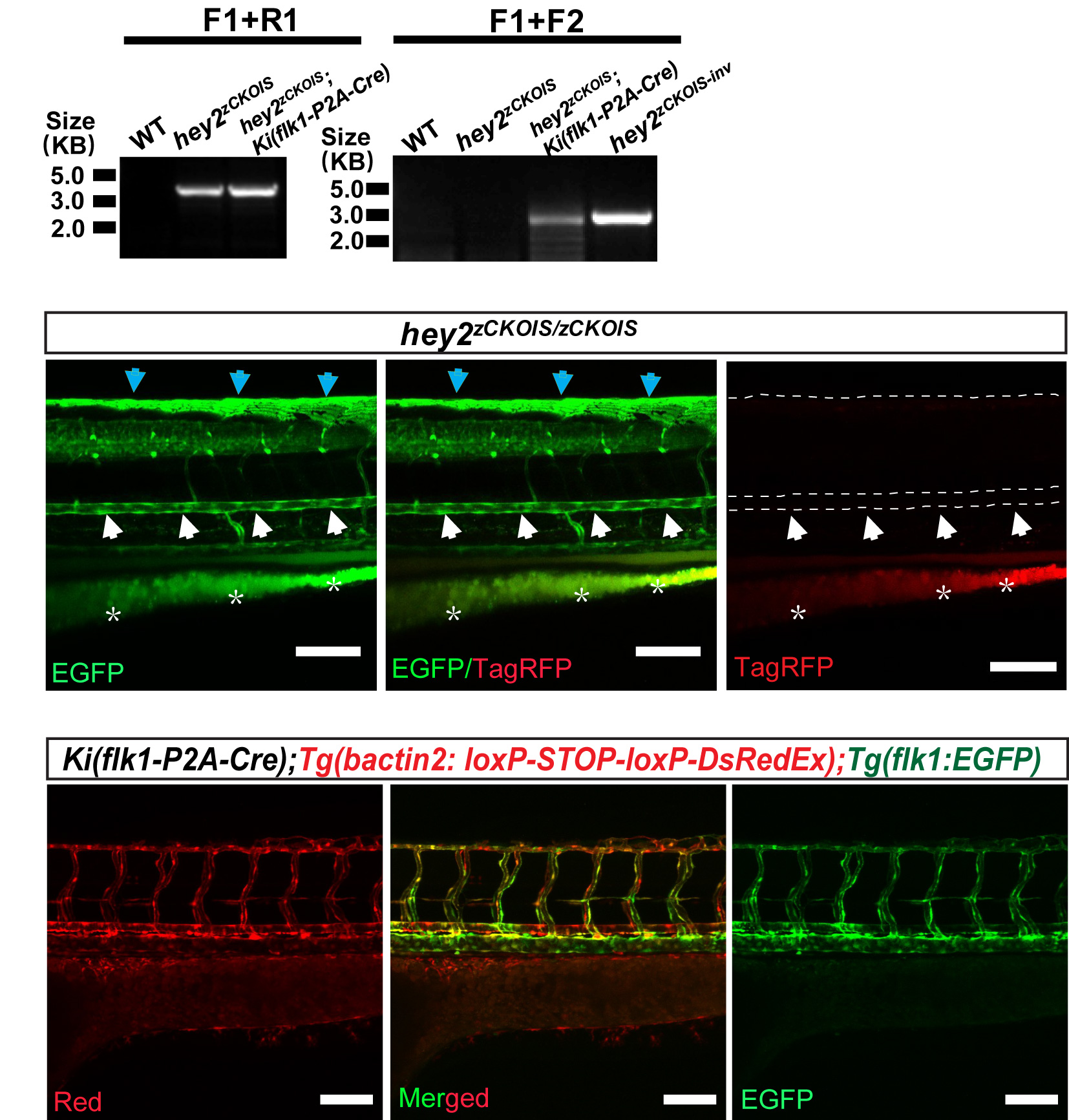
